# Evidence for positive priming of leaf litter decomposition by contact with eutrophic pond sediments

**DOI:** 10.1101/749101

**Authors:** Kenneth Fortino, Jessica Hoak, Matthew N Waters

## Abstract

Organic matter processing controls the flow of carbon and nutrients through ecosystems. Heterotrophic metabolism within ponds is supported by both terrestrial leaf litter and autochthonous production. We investigated the potential for the priming of leaf litter decomposition in small ponds using microcosms. We incubated senescent tulip poplar (Liriodendron tulipifera) leaf discs in the dark for 130 days either in contact with eutrophic pond sediments or isolated from sediment contact. Leaves that had been in contact with the sediments were significantly less tough and lost more carbon mass following the incubation than leaves that were not in contact with the sediments, indicating that they were decomposing faster. We calculated a positive priming effect of the sediments of 42% and 77% based on the change in toughness and C mass loss, respectively. We further found that leaf discs that were in contact with the sediments had significantly less fungal biomass, measured as ergosterol mass, and less leaf-derived N in fungal biomass than the leaf discs isolated from the sediments. These results indicate that the presence of the more labile organic matter of the sediments alters the rate of organic matter mineralization and the cycling of nitrogen and carbon.

## Introduction

Organic matter processing is a fundamental ecosystem function that regulates the flow of energy and matter through biogeochemical systems (Schlesinger and Bernhardt 2013). The availability of energy and inorganic nutrients to an ecosystem is controlled by the mineralization rate of accumulated organic matter by microbial and animal communities (Schlesinger and Bernhardt 2013). There is substantial spatial and temporal variation in the features that combine to affect organic matter mineralization rate in inland waters (i.e., oxygen concentration, temperature, inorganic nutrient concentration, and organic matter quality), making these systems highly dynamic venues of organic matter processing (Hargrave 1969, Granéli and Granéli 1978, del Giorgio and Cole 1998, Wetzel 2001, Berggren et al. 2010, Sobek et al. 2011, Fortino et al. 2014, Gudasz et al. 2015). Diversity in the source and quality of organic matter substrates is an important component of the observed variation in mineralization rate. Metabolism in most inland waters is supported by a combination of primary production within the system (i.e., autochthonous organic matter) and subsidies of allochthonous organic matter from the watershed (Wetzel 2001, Marcarelli et al. 2011). The mixture of autochthonous and allochthonous organic matter sources mean that the organic matter pool of a water body consists of a complex combination of sources, which vary in reactivity and quality (Meyers and Ishiwatari 1993, Finlay and Kendall 2008).

Considerable research has shown that the complexity of organic matter pools can alter mineralization in non-additive ways (Gartner and Cardon 2004). In lentic systems, the source and reactivity of dissolved organic matter sources was reported to alter system metabolism (Wetzel 1992, Kritzberg et al. 2004, Carpenter et al. 2005, Guillemette and del Giorgio 2011). In lotic systems, differences in the diversity of leaf species within mixed litter pools could increase (Swan et al. 2009) or decrease (Kominoski et al. 2007; Rosemond et al. 2010) the rate of litter decomposition. Moreover, the effect of organic matter diversity on decomposition is variable with environmental context (Lecerf et al. 2007) and can be altered due to nutrient enrichment (Rosemond et al. 2010), or season (Swan and Palmer 2004). Therefore, understanding the variation in organic matter processing and its impact on carbon and nutrient cycling, requires understanding the effect of complex organic matter source pools on mineralization.

One model describing the impact of interacting organic matter pools on mineralization is priming (Guenet et al. 2010; Bianchi 2011). Priming is a special case of organic matter interactions where the mineralization of refractory organic matter sources is stimulated by the addition of a more labile source of organic matter (Guenet et al. 2010; Bianchi 2011). Priming effects are commonly observed in terrestrial soils but remain poorly described in inland waters (Guenet et al. 2010, Bengtsson et al. 2018, Halvorson et al. 2019b). Investigations of priming in inland waters have found both evidence of positive priming and the absence of priming effects (Bengtsson et al. 2018, Halvorson et al. 2019b). However, the investigation of priming in inland waters was focused primarily on the stimulation of the breakdown of allochthonous dissolved organic carbon (DOC) via the addition of very simple organic substrates in lakes (Guenet et al. 2014; Bianchi et al. 2015; Dorado-García et al. 2015), and on periphyton and leaf litter interactions in streams (Halvorson et al. 2019b).

The emphasis on evaluating complex leaf litter pools in streams and complex DOC pools in lakes makes sense given that, in terms of system metabolism, these are the dominant forms of organic carbon in these systems, respectively (Wetzel 2001). However in some freshwater systems (e.g., small ponds), leaf litter decomposition occurs in close juxtaposition to water-column and benthic primary production, creating spatial overlap between the decomposition of allochthonous particulate organic matter sources (i.e., terrestrial leaf litter) and autochthonous organic matter sources (i.e., algal detritus). Organic exudates from living algae have been shown to prime microbial saprotrophs on leaf litter (Danger et al. 2013; Kuehn et al. 2014) but to our knowledge it has not been determined whether algal-derived detritus would have a similar priming effect.

Small ponds are very abundant (Downing et al. 2006, Hanson et al. 2007, Dowing 2010) and contain both high organic matter mineralization (Kortelainen et al. 2006, Tranvik et al. 2009) and storage (Tranvik et al. 2009), which can have a disproportionate effect on watershed-scale organic matter processing (Downing 2010). The majority of small ponds are expected to be eutrophic and have high watershed connectivity (Fairchild et al. 2005). As a result, small ponds should have high allochthonous and autochthonous organic matter loading, and contain an organic matter pool that is divergent in reactivity and quality, similar to what is found in forest soils. Thus, as with forest soils, we would predict that there would be a high potential for priming effects in the metabolism of the accumulated pond sediment organic matter (Bengtsson et al. 2018). The ubiquity of small ponds and their potential for complex organic matter and mineralization interactions (including priming) means that understanding the factors that affect the fate of the organic matter pool in ponds is essential to understanding watershed carbon cycling when small ponds are present.

For this study, we hypothesized that the presence of sediment organic matter from a eutrophic pond (i.e., partially labile autochthonous organic matter) would alter the decomposition of leaf litter subsidies (i.e., refractory, allochthonous organic matter). We tested this hypothesis using a microcosm system where we measured the decomposition of leaf discs in contact with sediments (and the labile organic matter pool), and separated from sediments. Leaf decomposition was measured as ash-free-dry-mass loss, change in leaf toughness, and change in C mass for leaves. To evaluate the role of fungal colonization on differences in litter decomposition rate, we also measured ergosterol mass on the leaves, following the incubation.

## Materials and Methods

### Experiment Set–up

The effect of sediment contact on the rate of leaf decomposition was tested in microcosms made from 250 ml (13 cm tall X 5 cm diameter) wide–mouth glass jars. We added known amounts of sediments and water to the jars from Lancer Park Pond (37.3062 N, −78.4043 W), a 0.1 ha eutrophic, constructed pond with a 1.5 m maximum depth. The sediments were collected from the pond on January 29, 2016 with an Ekman dredge at 2 different locations approximately 20 m from the shore. The collected sediments were homogenized and passed through a 250 μ m mesh net to remove macroinvertebrates and coarse particulate matter. We added 100 ml of the homogenized sediment slurry to each jar and then filled the remaining volume of the jar with water collected from the surface of the pond. We did not measure the pond water nutrient concentrations at the time of collection but summertime DIN and DIP concentrations are below 0.1 and 0.01 mg L^-1^ (K. Fortino, unpublished data). We determined the organic matter content and ash-free-dry-mass (AFDM) of the sediments with 3, 10 ml samples of this sediment and water mixture. The samples were dried at 50° C for 48 h and then ashed at 550° C for 5 h to determine loss on ignition (LOI).

To establish a natural sediment and water column environment we allowed the sediments to settle in the dark at room temperature over a 15-day period. During this time the overlying water was replaced twice by siphoning out the existing water and gently adding new water, taking care not to disturb the sediments. We replaced the overlying water on February 2, 2016 and February 5, 2016 with water that had been collected from the pond on January 29, 2016 and February 4, 2016, respectively. All pond water that was used for these replacements and to replace evaporative losses during the incubation was stored in the dark at 4° C.

The leaf litter we used in the microcosms was from a stock of tulip poplar (*Liriodendron tulipifera*) leaves that were collected at senescence in 2013 and then air dried. The leaves were stored dried in paper bags in the dark at room temperature before being used in the experiment. To prepare the leaf litter for the mesocosms, we soaked 35 leaves in 10 L of deionized water for 5 days to remove any immediately soluble organic matter. We then cut 13.5 mm diameter leaf discs from the leaves with a # 7 cork borer. These leaf discs were then soaked in deionized water for 2 days at 4° C to insure they would sink when added to the microcosms.

On February 12, 2016 we added 10 randomly chosen leaf discs to each microcosm. These leaf discs sunk to the bottom of microcosm and were in contact with the sediments for the duration of the incubation. Another 10 randomly chosen leaf discs were added to each microcosm on top of a 2.0 cm X 3.5 cm shelf made out of 0.5 cm mesh steel hardware cloth. These shelves were held 4 cm off of the sediment surface by 2 extensions of the hardware cloth shelf that were bent downward to form legs. An additional 9 replicates of 10 leaf discs were randomly selected and each dried at 50° C for 48 h and then ashed at 550° C for 5 h to determine the initial leaf organic matter content via loss on ignition.

Leaf samples for initial percent C and N were prepared by cutting 40, 10 mm leaf discs from the same tulip poplar litter that was used for the microcosm additions. Prior to cutting the leaf discs the litter was soaked for 72 h in deionized water in the dark at room temperature. The leaf discs were randomly divided into 2 samples of 20 discs each and then analyzed for C and N as described below.

### Leaf break–down and sampling

The microcosms were incubated for 130 days in the dark at room temperature. Volume lost to evaporation was replaced approximately weekly using pond water that was collected on February 10, 2016 and stored at 4° C. At the conclusion of the incubation the overlying water was siphoned out of each microcosm and we collected the following samples from both the leaves in contact with the sediments and on the shelf: two discs were preserved in 10 ml of HPLC-grade methanol and stored in at −20° C for ergosterol analysis, three discs were gently dried with a paper towel and used to determine leaf toughness and afterward, C:N. The remaining 5 leaf discs were dried and ashed as described above to determine AFDM via LOI. All of the leaf discs except those used to measure ergosterol were gently rinsed with deionized water before processing further.

### Ergosterol content and C:N ratio

The ergosterol content of the leaf discs was determined by first saponifying the samples at 80° C for 30 minutes in 10 ml of 8 g L^-1^ potassium hydroxide in methanol. Following this, the samples were allowed to cool and 3 ml of deionized water was added to each sample. The samples were then rinsed 3 times with 10 ml of pentane followed by 30 seconds of vortexing. Following each rinse, the pentane layer was removed and evaporated under N_2_. Once all the pentane had been evaporated, the pellet was resuspended in 1 ml of methanol and ergosterol mass was determined using a Shimadzu 10VP HPLC equipped with a Whatman Partisphere C18 reverse-phase column set at 40 degrees C. The UV detector was set to 282 nm and a 100% methanol flow at 1 mL per minute.

Leaf toughness was determined with a penetrometer (Graça and Zimmer, 2005) that used a 2 mm diameter cylindrical punch attached below a reservoir. Each leaf disc was placed under the punch and water was slowly added to the reservoir until the punch penetrated the leaf disc. After being used in the penetrometer, each leaf disc was dried at 50° C for 48 hours and ground using a stainless steel spatula. Carbon and nitrogen content (as percent) were then measured on the ground leaf material using a Costech C-H-N Combustion Analyzer. Samples were acidified in HCl vapors for 24 hours prior to analysis to remove inorganic carbon. Sample C:N was determined as the molar ratio of C and N in the sample.

### Calculations and statistical analysis

The AFDM of the leaf discs was calculated as LOI after ashing at 550° C. The AFDM of a single leaf disc was estimated as the AFDM of the total sample divided by the number of leaf discs in the sample. Leaf mass loss was calculated as the difference between the estimated initial AFDM mass of a single leaf disc prior to the incubation and the estimated final AFDM mass of a single leaf disc from each position after the incubation.

Since 3 leaf discs were subsampled to measure toughness from each position in each bottle, we averaged the mass required to puncture each of these leaf discs to get a single mass for each bottle. This average mass was used as the measure of leaf toughness in the analysis.

To be able to compare our results to other studies that measured priming using different methods, we estimated the priming effect as a percentage of the amount of leaf decomposition based on either the toughness or the C mass loss of the leaf discs not in contact with the sediments (Bengtsson et al. 2018). We calculated the priming effect as PE = (R_p_/R_c_) * 100 where PE is the priming effect as a percent, R_p_ is the difference between the response of the leaf discs in contact with the sediments and those not in contact with the sediments, and R_c_ is the response of the leaf discs not in contact with the sediments. Since greater decomposition results in a lower measurement value of toughness, the PE based on toughness would be negative when there is positive priming. To make this result consistent with the standards of the literature (i.e., positive PE = positive priming), we report PE based on toughness with the sign reversed.

To estimate the changes in C and N mass following the incubation, we estimated the C and N mass of a single leaf disc as the estimated AFDM of a single leaf disc (see above) multiplied by the proportion of C or N. To calculate the percent contribution of fungal carbon or nitrogen to the carbon or nitrogen mass of a leaf disc at the end of the incubation, we converted ergosterol mass to fungal mass assuming that there is 1 mg of fungal dry mass per 5 μ g of ergosterol (Su et al. 2015). The carbon and nitrogen mass in the fungal biomass was determined by assuming that 43% and 6.5% of fungal dry mass was carbon and nitrogen, respectively (Findlay et al. 2002). The percent contribution of fungal carbon or nitrogen mass to the carbon or nitrogen mass of the leaves following the incubation was calculated as the fungal carbon or nitrogen mass divided by the leaf carbon or nitrogen mass, then multiplied by 100.

To assess the degree to which fungal nitrogen immobilization could account for the reduction in leaf C:N following incubation, we first determined the mass of nitrogen that would have been lost from the leaves had the C:N not changed during the mass loss associated with decomposition. In other words, if carbon and nitrogen had been lost from the leaves during decomposition in the same ratio that they were found in the pre-incubation leaf tissue. The estimate of the leaf nitrogen mass following the incubation without immobilization was calculated by dividing the leaf carbon mass following the incubation by the mass-based C:N. The average of this estimated final leaf nitrogen mass was then subtracted from the initial nitrogen mass of the leaves to estimate the total nitrogen mass that was mineralized during decomposition. The proportion of mineralized nitrogen that was potentially immobilized by the fungi was then estimated as the average fungal nitrogen mass divided by the estimate of the total mass of nitrogen mineralized from the leaves.

Since the leaf discs on the shelf (i.e., not in contact with the sediments) and those on the sediments of a single bottle cannot be assumed to be independent, we analyzed the all differences between leaf discs in contact or not in contact with the sediments by calculating the difference in estimated mass loss (based on AFDM, C mass, or, N mass) of a single leaf disc on the shelf from the estimated mass loss of a single leaf on the sediments. We then used a one sample t-test to determine if the difference in mass loss was significantly different from 0. All calculations and statistical analyses were performed using R (R Core Team 2014).

## Results

The mean (± 1 SD) dry mass of sediment in the microcosms was 8.3 (± 0.11) g. The sediment averaged 16.8 % organic matter and thus contained 1.4 g (dry mass) of organic matter. A single leaf disc added to the microcoms had a mean (± 1 SD) AFDM of 0.0035 (± 0.0004) g, so the initial AFDM of 20 leaves added to each microcosm provided approximately 0.071 g of organic matter to the microcosms.

Contact with the sediments accelerated the decomposition of the leaf discs compared to those that were incubated above the sediment-water interface. The leaf discs that were not in contact with the sediments were significantly more tough, requiring an average of 16.4 g more weight to puncture the leaf disc than those that were in contact with the sediments (t = 4.033, p = 0.00296; Figure 1). The loss of toughness in the leaf discs occurred concomitantly with the loss of carbon from the leaf tissues (Table 1, Figure 1). A leaf disc in contact with the sediments lost 0.304 mg more C mass (t = −2.798, p = 0.0207; Table 1) and had 8 percent less C than a leaf disc not in contact with the sediments (t = 8.403, p < 0.0001; Table 1, Figure 1).

**Table 1.** The mean (± 1 SD) AFDM, C mass, and N mass of a single leaf disc (mg), and the percent C, percent, N and C:N of the leaf discs before incubation and after incubation in contact with pond sediments or not in contact with pond sediments. The initial AFDM was calculated from 9 replicate samples of 10 leaf discs. The AFDM of each replicate was divided by the number of leaf discs in the replicate to estimate the AFDM of a single leaf disc. The AFDM of a leaf disc following incubation was as determined in the same way as for the initial sample except that each treatment level had 10 replicate samples of between 4 and 6 leaf discs. The percent C and N of the leaf discs prior to incubation (Initial) was determined from 2 replicate samples of 20 leaf discs, and the percent C and N of the leaf discs following incubation was determined from 10 replicate samples of 3 leaf discs from each treatment level. In all cases C:N is calculated as the molar ratio.

| Source | AFDM | Percent C | C Mass | Percent N | N Mass | C:N |
| --- | --- | --- | --- | --- | --- | --- |
| Initial | 3.54<br>(± 0.42) | 45.13<br>(± 0.94) | 1.60<br>(± 0.19) | 0.99<br>(± 0.06) | 0.035<br>(± 0.004) | 53.43<br>(± 4.30) |
| Sediment<br>Contact | 2.49<br>(± 0.43) | 36.90<br>(± 3.06) | 0.90<br>(± 0.16) | 1.58<br>(± 0.16) | 0.038<br>(± 0.007) | 27.27<br>(± 1.40) |
| No Sediment<br>Contact | 2.68<br>(± 0.54) | 45.02<br>(± 0.85) | 1.20<br>(± 0.23) | 1.57<br>(± 0.20) | 0.043<br>(± 0.013) | 34.06<br>(± 4.73) |

**Figure 1.**
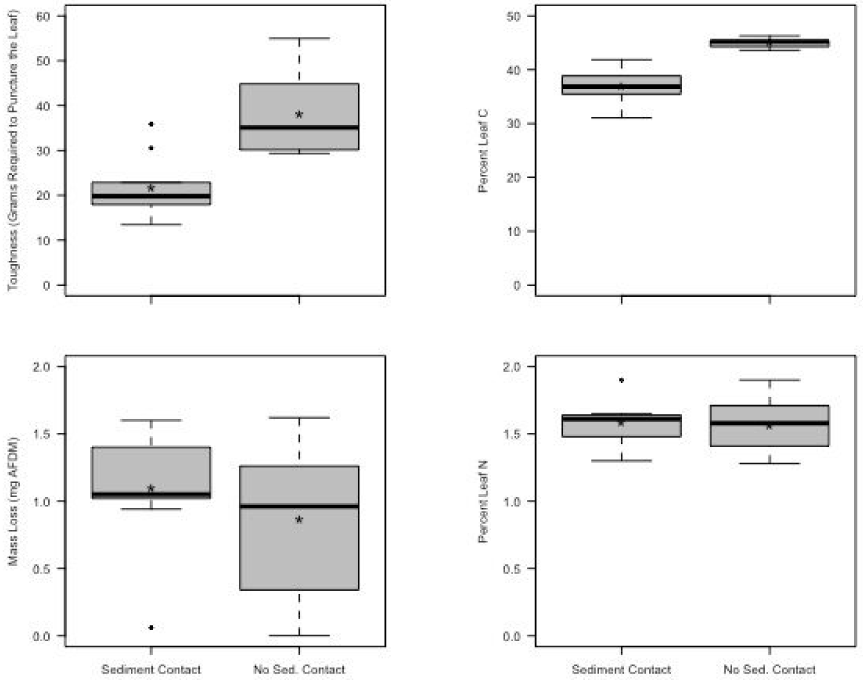
The mean mass required to puncture the leaf discs (toughness), percent leaf C, percent leaf N, and mass loss estimated AFDM lost from a single leaf of the leaf discs, following incubation in contact, or not in contact with the sediments in the microcosm. Each boxplot shows the results from the 10 replicate microcosms where, the horizontal line indicates the median mass required to puncture the leaf, the box encloses the 1^st^ and 3^rd^ quartiles, the whiskers indicates the range, and the mean is shown by the star. Individual points indicate outlier observations.

Despite the increased loss of toughness and C from the leaves in contact with the sediments, we did not detect a difference in the AFDM loss of the leaves in contact with the sediments and not in contact with the sediments. The mean (± 1 SD) AFDM loss of the leaf discs in contact with the sediments was 1.101 (± 0.432) mg (32% of the original mass), and the AFDM loss of the leaves not in contact with the sediments was 0.864 (± 0.536) mg (25% of the original mass) but the difference between the loss of AFDM from the different treatments was not significantly different from 0 (t = −0.857, p = 0.4136; Figure 1). The PE based on leaf toughness (42%) and the PE based on C mass loss (77%) showed evidence of a substantial positive priming effect of sediment contact on the decomposition of the leaf litter.

In contrast to C, there was no effect of sediment contact on the change in N mass or percent N in the leaf. A leaf disc gained a mean (± 1 SD) of 0.006 (± 0.0103) mg of N (Table 1) during the incubation but the difference in the change in N mass (t = −0.725, p = 0.487, Table 1, Figure 1) or percent N (t = −0.183, p = 0.859, Table 1) between leaves not in contact, and in contact with the sediments was not significantly different from 0. The effect of the sediment contact on C content but not N content of the leaf discs unsurprisingly resulted in a significant difference between the C:N of the leaves in contact with and not in contact with the sediments (t = 3.963, p = 0.0033; Table 1). In both treatment levels, the C:N of the leaf discs declined from the initial ratio during the incubation but the leaf discs in contact with the sediments lost more C than those not in contact with the sediments, resulting in a greater decline in C:N (Table 1).

When not in contact with the sediments, the leaf discs had 173 μg (g AFDM)^-1^ more ergosterol (t = 5.2436, p = 0.0005) than the leaves in contact with the sediments, which indicates a difference of 34.6 mg (g AFDM)^-1^ more fungal biomass on the leaves not in contact with the sediments (Table 2). The greater fungal biomass on the leaves not in contact with the sediments meant that fungal C and N made up a significantly greater proportion of the total C (t = 3.553, p = 0.0062) and N (t = 5.183, p = 0.0006) mass of the leaves not in contact with the sediments following the incubation (Table 2).

**Table 2.** The mean (± 1 SD) of the ergosterol mass normalized to leaf AFDM (μg (g AFDM)^-1^). Mean (± 1 SD) fungal biomass (mg (g AFDM)^-1^) estimated from the ergosterol mass using 1 mg of fungal biomass per 5 μg of ergosterol, the fungal carbon mass and fungal nitrogen mass of a single leaf disc (mg (g AFDM)^-1^) estimated using fungal C and N percentages of 43% C and 6.5% N. The Mineralized N Mass (mg) is the estimated mass of N lost from the leaves due to mineralization and the Percent Fungal N Immobilization is the percent of the mineralized N mass that can be accounted for by fungal N mass.

| Source | Ergosterol Mass | Fungal Biomass | Fungal C Mass | Fungal N Mass | Mineralized N Mass | Fungal N Immobilization |
| --- | --- | --- | --- | --- | --- | --- |
| Sediment Contact | 119.8<br>($\pm 71.9$) | 24.0<br>($\pm 14.4$) | 10.3<br>($\pm 6.19$) | 1.56<br>($\pm 0.935$) | 0.0153 | 24.2 |
| No Sediment Contact | 292.7<br>($\pm 78.9$) | 58.5<br>($\pm 15.8$) | 25.2<br>($\pm 6.78$) | 3.81<br>($\pm 1.03$) | 0.0086 | 120.8 |

Using the above information, we developed C and N mass balance models for when the leaves are, or are not in contact with the sediments (Figure 2). These models show that when the leaf discs are not in contact with the sediment 0.069 mg of the 0.40 mg of C lost from a leaf disc are found in fungal biomass, leaving 0.33 mg or 83% of the C lost from the leaf due to other microbial and mineralization processes. When the leaf discs are in contact with the sediments, they lost more C mass (Figure 1, Table 1) but due to the lower fungal biomass on the leaf discs, only 0.024 mg C are in the fungal biomass, leaving 0.68 mg or 97% of the C lost from the leaves due to other microbial and mineralization processes.

**Figure 2.**
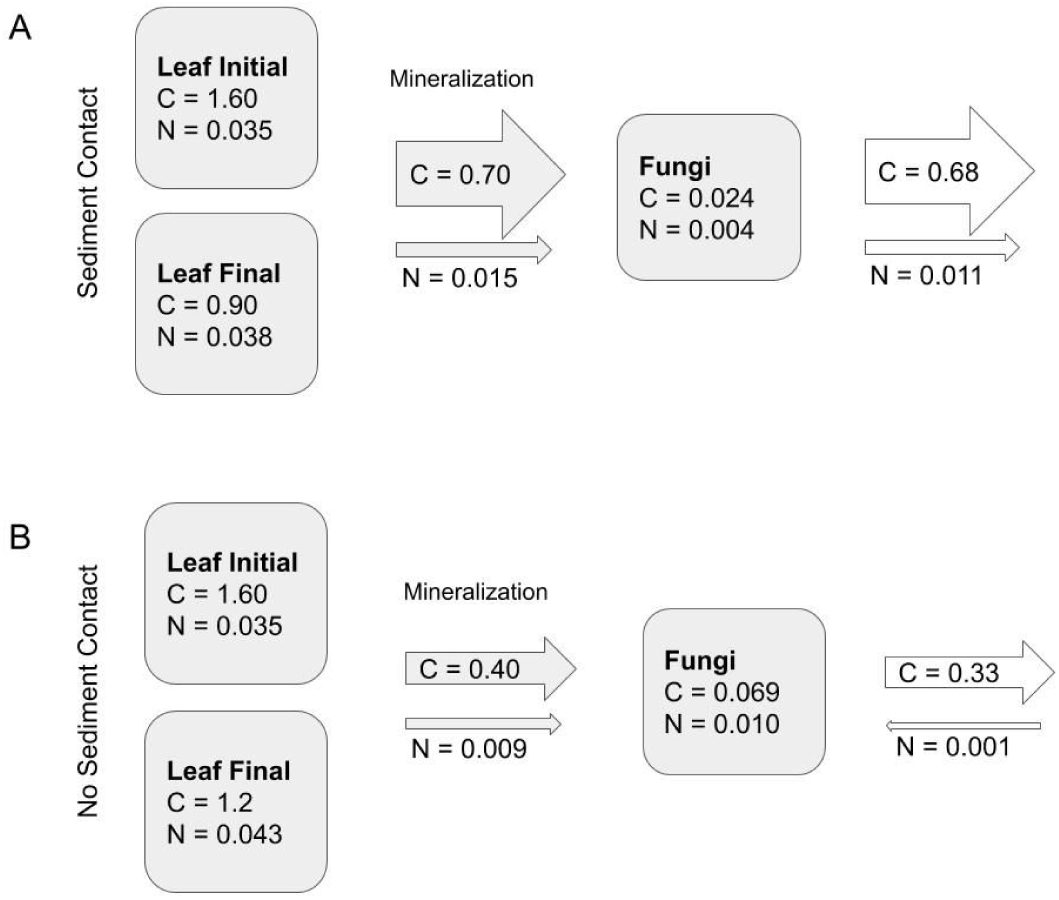
A carbon and nitrogen mass balance model for the decomposition of the leaf discs in contact with the sediments (A) and not in contact with the sediments (B). The model shows the amount of C and N mass (mg) estimated for a single leaf disc at the beginning of the incubation (Leaf Initial), at the end of the incubation (Leaf Final), and in fungal biomass at the end of the incubation (Fungi). The mass (mg) of C and N lost from the leaves during the incubation and therefore available for fungal uptake is shown in the arrow labeled “Mineralization”. The white arrows facing away from the fungi show the mass of C or N that was not required for fungal biomass and would be either exported or available for other consumers (e.g., bacteria). The white arrow facing toward the fungi shows the N mass that would need to be immobilized to meet fungal demand.

Assuming that N mass was lost from the leaves due to mineralization proportional to the C:N of the leaf, we estimated that during the incubation a single leaf disc would have lost 0.0153 mg of N when in contact with the sediments and 0.0086 mg of N when not in contact with the sediments (Figure 2, Table 2). When not in contact with the sediments, the fungal biomass on a single leaf contained an estimated 0.01 mg of N, which is greater than the N released from a leaf disc by mineralization, and would require the immobilization of 0.001 mg of exogenous inorganic N (Figure 2). Because of the lower fungal biomass on the leaf discs that were in contact with the sediments, there was only 0.004 mg of N in fungal biomass at the end of the experiment (Table 2) which represents only 26% of the N mineralized from the leaf disc (Figure 2, Table 2).

## Discussion

### Priming of leaf litter decomposition

The results of our experiments show that leaf litter decomposition is accelerated by contact with the sediments of a small eutrophic pond and suggest that organic matter processing rates are affected by the complexity of the available organic matter. Although our experiment cannot elucidate the specific mechanism affecting the leaf litter decomposition rate, our findings are consistent with priming. Priming is an increase in the mass loss from the refractory organic matter pool as a result of the addition of a labile organic matter (Guenet et al. 2010; Bianchi 2011). In our microcosms, the leaf litter was refractory relative to the sediment organic matter, based on the C:N of the different organic matter pools.

Differences in C:N reflect differences in organic matter source and reactivity (Ostrofsky 1997, Kaushal and Binford 1999, Wetzel 2001). Autochthonous organic matter derived from phytoplankton production is less carbon rich due to the fact that algal biomass typically does not contain the complex structural carbohydrates and lignins found in vascular plants (Meyers 1994) and typically has a C:N of less than 12 (Meyers 1994, Wetzel 2001). Thus, the C:N of 10 observed for the pond surface sediments in this system (K. Fortino, unpublished data) suggested that they are derived primarily from autochthonous detritus.

In contrast to the pond sediments, the *L. tulipifera* leaf litter used in the microcosms had an average C:N of 53 prior to incubation, which is slightly lower than what has been observed for “typical” terrestrial leaf litter C:N (Aerts 1997; McGroddy et al. 2004). This recalcitrant material would be stoichiometrically out of balance with microbial consumers and therefore be nutritionally limiting and require that they immobilize exogenous nutrients to maintain growth (Suberkropp and Chauvet 1995).

Although the change in AFDM did not reveal a significant difference in the overall mass lost from the leaves in the different conditions, the leaves in contact with the sediments were less tough and lost more C than the leaves that were not in contact with the sediments. Leaf toughness results from the structural carbohydrates found in leaf tissue (Gessner 2005) and the loss of toughness is correlated with the loss of mass via decomposition (Grubbs and Cummins 1994, Medeiros et al. 2009). The reduced C content of the leaves in contact with the sediments also provides evidence of increased leaf decomposition in the presence of the sediment organic matter, since carbon is the major structural element in plant tissue (Schlesinger and Bernhardt 2013). Taken together, these results indicated that the refractory organic matter of the leaves was mineralized faster when in contact with the labile organic matter in the sediments in a way that is consistent with priming.

### Magnitude of Priming

We measured a positive priming effect of the sediment environment on the decomposition of the leaf litter when we evaluated decomposition based on the change in toughness and the change in carbon mass. Since we measured no significant difference in the change in AFDM between the leaf discs in contact, and not in contact with the sediments, we found no priming effect when decomposition was measured as AFDM loss.

The total percent mass lost in our experiment (25 - 32% of original mass) was somewhat lower than the 40 - 60% mass loss that has been observed in other studies (Danger et al. 2013, Soares et al. 2017). This relatively low mass loss overall combined with the background variation that results from the limitations of measuring changes in mass from low mass leaf samples may have reduced our ability to detect the relatively small variation in leaf mass loss due to priming.

The priming effect based on carbon mass change (77%) was greater than the priming effect based on the change in leaf toughness (43%) but both measures were substantially greater than the mean 12.6% priming effect for aquatic systems reported by Bengtsson et al. (2018). However, Bengtsson et al. (2018) found that, overall aquatic systems show highly variable and not consistently positive priming effects. Furthermore, they hypothesize that priming effects will predominantly be observed in sediment environments, where they calculated a mean priming effect of 46.2% for marine sediments (Bengtsson et al. 2018) that was similar to what we observed for the leaf litter in the pond sediments. The composition of our test sediments was primarily composed of labile autochthonous organic material (∼ C/N 10), which supports our priming numbers being greater than other systems that contain more structural carbon in the sediments.

The magnitude of the priming effect that we observed for the pond sediments on the leaf litter was similar to other examples of positive priming of leaf litter decomposition. Danger et al. (2013) found a priming effect for algae additions to stream-derived leaf litter of approximately 43%, which is similar to what we found for the PE based on the change in toughness of leaves that were in contact with pond sediments. In contrast, Halvorson et al. (2016), found a much higher priming effect of approximately 167% when algae were allowed to grow on decomposing litter without P additions. Overall, the magnitude of the priming effect that we observed was consistent with the, albeit highly variable and limited, positive priming effects recorded in the literature under ambient nutrient conditions.

### Effect of sediment contact on nitrogen cycling

Danger et al. (2013) and Halvorson et al. (2016) only found positive priming effects under conditions of nutrient limitation. The nutrient status of our microcosms is unknown, but our observation that the leaf discs lost C mass, without a simultaneous loss of N mass, suggested that the microbial community was immobilizing inorganic nitrogen liberated during decomposition (Suberkropp and Chauvet 1995) and suggested microbial N limitation. The select mineralization of C and immobilization of N is an established pattern in leaf conditioning by microbial communities (Anderson and Sedell 1979) and was reflected in the decrease in C:N of the leaf tissue during our experiment.

Interestingly, the presence of the labile organic matter in the sediments appears to alter the way that N is cycled in the leaf-decomposer system. When labile sediment organic matter was not present, the mass of inorganic nitrogen liberated from the leaves via mineralization was insufficient to support the growth of the observed fungal biomass, and the fungi must have immobilized inorganic nitrogen from sources other than the leaf tissue (Figure 2). In contrast, when the leaves were in contact with the labile sediment organic matter, the combination or greater leaf mineralization and lower fungal biomass meant that there was more inorganic nitrogen released from the leaf tissue than was required for the observed fungal biomass and nitrogen would have been lost from the system (Figure 2).

The implications of the differences in N cycling in the presence or absence of sediment contact are that the sediment contact may influence the nitrogen limitation of the bacterial decomposers on the leaf tissue. If we assume that the only fungal biomass production was what we measured at the conclusion of the incubation and that fungi utilize all of the available nitrogen liberated from the leaves before the bacteria, then there would be no leaf-derived N available for the bacteria in the absence of the sediments but 73% of the leaf-derived nitrogen in the presence of the sediments. Obviously these assumptions are invalid due to fungal biomass losses and real-time competition for N by the bacteria (Gessner and Chauvet 1994, Gessner 1997, Baldy et al. 2002, Halvorson et al. 2019b), but the measurements suggested that the sediments may alter the N availability to the microbial community on the leaves.

### Microbial Dynamics

The priming effect observed in our experiment was likely the result of a change in microbial decomposition dynamics, since larger metazoans were excluded from the mesocosms. Despite meiofaunal burrowing activity that could have occurred in some of the mesocosms, we do not believe that this contributed to the differences in decomposition. Patterns of microbial organic matter colonization have been well studied in streams, and in these systems fungal biomass and production are greater than bacterial biomass and production (Weyers and Suberkropp 1996, Baldy et al. 2002) and fungi typically dominate leaf litter mass losses (Gessner and Chauvet 1994). In streams with complex organic substrates, there is some partitioning of the substrate where fungi are more abundant and active on coarse substrates and bacteria are more active on fine organic matter substrates, although fungi are still typically more abundant overall (Gessner 1997, Baldy et al. 2002, Findlay et al. 2002, Findlay 2010).

Temporal dynamics of fungal biomass on leaf litter are variable but fungal biomass on leaf litter in lotic systems has been found to reach a maximum of 50 to >100 mg (g leaf)^-1^ between approximately 20 to 40 days of incubation and then decline (Baldy et al. 1995, Weyers and Suberkropp 1996, Baldy et al. 2002, Pascoal and Cassio 2004, Danger et al. 2013). We measured between 24 and 58 mg (g AFDM)^-l^ of fungal biomass on the leaves after 130 days of incubation, which was likely after peak fungal biomass. Though comparable to the 40 mg C (g leaf C)^-1^ measured on leaf litter from a large river at 140 days (Pascoal and Cassio 2004), it was lower than the 7.5 g (g leaf AFDM)^-1^ fungal standing stock measured from a small stream (Findlay et al. 2002).

In our experiment, the leaves that were in contact with the sediments had lower fungal biomass after 130 days of incubation than the leaves that were not in contact with the sediments despite greater carbon loss from the leaves, thus the increased carbon mineralization on the leaves in contact with the sediments was not simply a function of increase in fungal biomass.

Since we do not have a measure of fungal production, we cannot determine if the reduced fungal biomass resulted in reduced fungal production. It is possible that increased fungal production was not observed as biomass due to losses via conidia production (Halvorson et al. 2019a).

Alternatively, the sediment environment may have been stressful for fungi and limited growth. Environments with low oxygen (Medeiros et al. 2009) and with high nutrients, low oxygen, and low flow (Pascola and Cassio 2004) showed reduced fungal biomass. The leaf discs in contact with the sediments may have had lower fungal biomass because the sediment environmental conditions (i.e., low oxygen, low flow, high nutrients) were suboptimal for fungal growth.

## Conclusions

Our results show that the environmental conditions associated with leaf decomposition can alter the processing of organic matter in aquatic systems. In particular we found that the presence of a labile organic matter pool (i.e., sediments) can accelerate the loss of C from terrestrial subsidies to aquatic systems. We propose that small ponds could represent a unique environment for organic matter processing within the river network due in part to their creation of a complex organic matter pool derived from phytoplankton detritus and terrestrial leaf litter. Our results suggest mechanisms of organic matter processing operating in lentic environments that are distinct from those in nearby lotic portions of the watershed.

## Author contributions

KF and JH designed and executed the study. KF analyzed and interpreted the data and wrote the manuscript. MW measured the C:N of the samples and provided feedback on the manuscript and analysis and interpretation of the data.

## Data Availability

The datasets generated during and/or analysed during the current study are available in the Zenodo repository, https://doi.org/10.5281/zenodo.2875782

## Acknowledgements

We thank Julia Marcellus and Jen Andrews for assistance with the field and lab work. We also thank Hal Halvorson and Kevin Kuehn for completing the ergosterol measurements. The manuscript was greatly improved by comments by Hal Halvorson and 2 anonymous reviewers. This study was partially supported by the Longwood University PRISM program.

## Notes

https://doi.org/10.5281/zenodo.2875782

